# Data-Driven Discovery of Feedback Mechanisms in Acute Myeloid Leukaemia: Alternatives to classical models using Deep Nonlinear Mixed Effect modeling and Symbolic Regression

**DOI:** 10.1101/2024.06.17.599366

**Authors:** Carl Julius Martensen, Niklas Korsbo, Vijay Ivaturi, Sebastian Sager

## Abstract

In pharmacometrics, developing and selecting models is crucial for quantitatively assessing drug-biological interactions, treatment planning, and gaining insights into underlying processes.

These validated models are essential for predictive analytics and strategic decision-making in drug development and clinical practice. Unlike traditional methods, machine learning (ML) offers a data-driven alternative to conventional, first-principle approaches. This paper presents an automatic method to derive unknown or uncertain (sub)models using longitudinal, heterogeneous data. Initially, we employ deep nonlinear mixed effect (DeepNLME) to train a neural network as a universal approximator, which then generates a parameterized representation of the underlying process. Subsequently, we apply symbolic regression to identify a set of potential models expressed as equations. Within the study parameters, the proposed method outperforms the baseline models and demonstrates the validity of both the DeepNLME approach and the symbolic regression approach.

## 1 Introduction

Developing models is an intricate and lengthy process [2]. It requires proficiency in the model’s specific domain and in mathematics. Additionally, programming skills are required for algorithm development and usage to ensure reproducibility of obtained results [3], [4]. A deep understanding of the basic mechanisms and their related mathematical representations, often expressed as a set of ordinary differential equations (ODE), is crucial. However, within a particular context, typically dominated by real-world data, there are many published models that reflect different assumptions about the underlying mechanisms. Depending on the data and the identifiability of the models, some parameters, such as reaction rates, must be sourced from an extensive review of the literature and often have a major effect on the final outcome of the model selection process. In pharmacometrics, the development and selection of models is essential for the quantitative assessment of drug-biological interactions. This encompasses the absorption, distribution, metabolism, and excretion of drugs within the body (pharmacokinetics) and their impact on the subject’s intrinsic metabolism (pharmacodynamics). Common applications of these models include the design of preclinical test campaigns, simulation studies on the effects of drugs [5], [6], or the investigation and derivation of (optimal) treatment schedules [7]–[9]. These rigorously validated models are crucial for predictive analytics and strategic decision making in drug development and clinical practice. Unlike traditional methods, machine learning (ML) offers a data-driven alternative to conventional first-principle approaches. Data is used to train universal approximators, which are a parameterized representation of the underlying process. This results in an algorithmic approach to constructing black-box models. Although many ML applications are used for everyday tasks, such as search engines or vision models, its application in scientific domains has only recently resurfaced as scientific machine learning (SciML) [10]. Numerous methods leverage universal approximators in scientific fields, as seen in [11]–[13]. These methods incorporate scientific assumptions about the structure or characteristics of the process at hand to seamlessly integrate a universal approximator into a rigorously defined model. Such grey-box models allow researchers to effectively detect deeply nested signals within the model, utilizing automatic differentiation and differentiable programming to optimize the parameters of the embedded approximator, ultimately producing data-driven oracles within the scope of the training data. To improve interpretability and potentially improve the extrapolation capabilities of the models, the collected data can be converted into a mathematical expression through symbolic regression, as demonstrated by [11], [14].

The application of SciML in the pharmaceutical field has seen substantial advancements in recent years. The use of ML with longitudinal data was explored in [15]. In [16], the authors applied ML to differentiate placebo responders in the context of eating disorders. The study in [17] used ML to select appropriate patients to curb the spread of COVID-19 with limited resources based on patient characteristics. The use of ML in survival analysis was examined in [18]. The impact and application of machine learning to optimize treatment schedules were investigated in [19]–[21]. In [22], the authors employed deep learning to deduce the parameters of a PK model from the chemical structure of the drug. The enhancement or learning of PK models has been studied in various works, such as [23]–[26]. In contrast, the use of symbolic regression to improve interpretability is much less common. A preprint [27] employs SR to identify relationships from data in the context of Preeclampsia. A more recent paper [28] uses neural ODEs to learn the dynamics of the model first and then derive an expression by visually inspecting the dynamics of the state space.

We present a fully data-driven method to derive unknown or uncertain (sub)models in the presence of longitudinal, heterogeneous data, as depicted in fig. 1. Opposing to the previously mentioned approaches to use symbolic regression, we a) provide a general workflow similar to [11] that benefits from scientific priors and b) enables the use of state-of-the-art symbolic regression software under the assumption of a shared model structure. To validate our method, we employ the procedure to develop a data-driven substitute for the traditional Fridberg feedback mechanism [29]. This mechanism describes the impact of maturing white blood cells in various models of leukemia. We refer to [1], which provides an in-depth analysis of different mathematical models of acute myeloid leukemia treated with high-dose cytarabine, along with the corresponding dataset.

**Figure 1.**
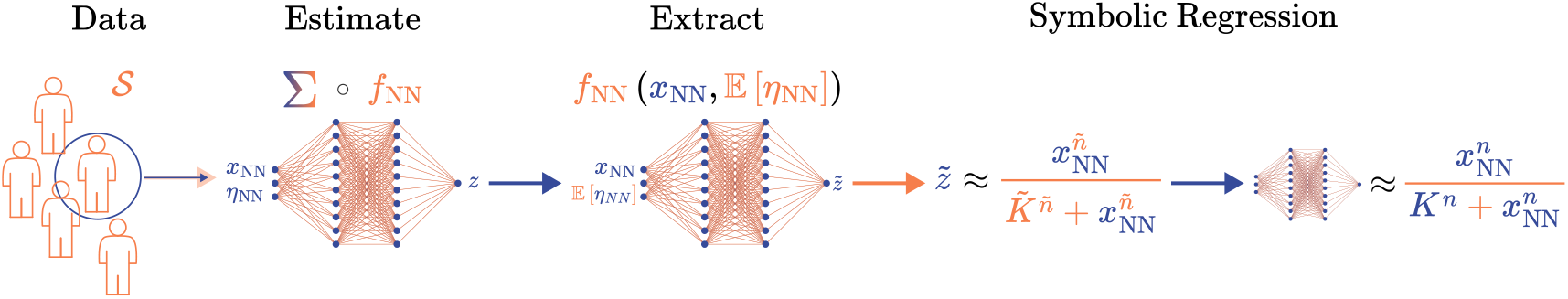
Graphical representation of the proposed workflow. The DeepNLME model Σ consists of a shared structure with an embedded neural network *f*_*NN*_ with a set of shared weights and biases and individual inputs. After training we use the typical values of the random effects and extract the networks typical response 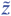 for all individuals in the population to perform symbolic regression. Finally the set of recovered functions, which are assumed to be shared among all subjects, can be used to perform model selection guided by an expert researchers domain knowledge. The final selection step can be performed by fitting either on the collected signal *z* or fitting the full model ∑.

**Figure 2.**
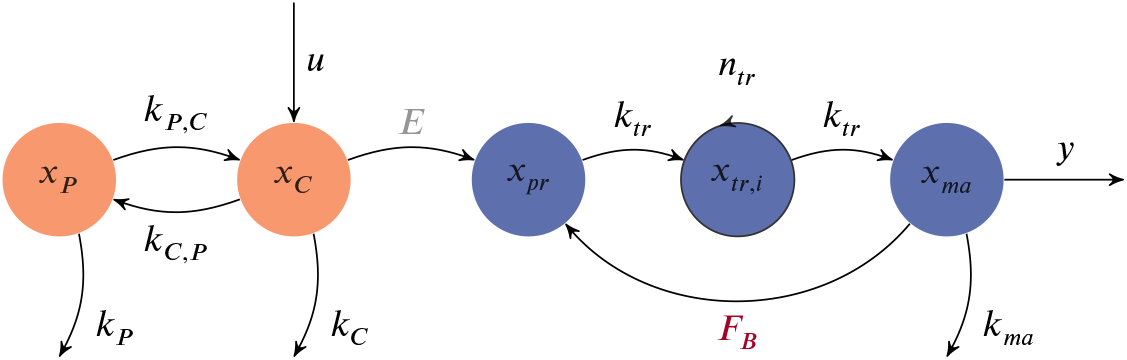
Graphical depiction of the dynamical model Equation (5). The pharmacokinetics is modeled using a Central-Peripheral system. The pharmacodynamic consists of the proliferation *x*_*pr*_ followed by *n*_*tr*_ transition compartments *x*_*tr*_. The maturating cells *x*_*ma*_ act on the proliferation via the feedback mechanism *F*_*B*_ which is represented by the neural network embedded in the dynamics Equation (5). For reasons of brevity, we define 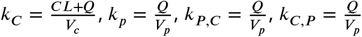.

The structure of the paper is as follows: Section 2 offers an initial overview of nonlinear mixedeffect models, symbolic regression, and a summary of the data set. Section 3 presents the essential details on deep nonlinear mixed-effect (DeepNLME) models, the mechanistic baseline model, and our proposed method. Section 4 outlines the experiments performed, and section 5 discusses their results.

## 2 Background

### 2.1 Nonlinear Mixed-Effect Models

Traditionally, the creation of models for real-world applications has depended on extensive and often protracted research. This includes a broad spectrum of scientific investigations, as well as the design and implementation of appropriate experiments to confirm hypotheses. Fundamental to most models is a mathematical representation of the underlying process, which also demands from the modeler a profound knowledge of prevalent models, essential optimization techniques, and data handling before and after processing. This holds particularly true in the field of PKPD models, where data is typically heterogeneous, noisy, and sparse across all subjects, incorporating not just dosing schedules and observations, but also a variety of subject-specific covariates such as age, gender, and weight. In mathematical terms, this is generally formulated using nonlinear mixed effect models (NLME) with the corresponding optimization problem across all subjects *i* from a specified population 𝒮

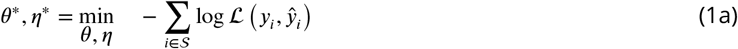

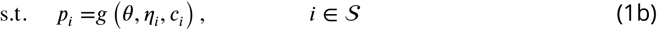

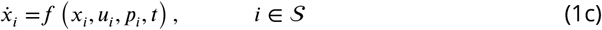

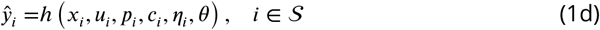

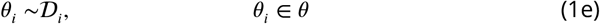

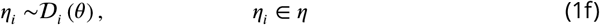

Where θ ∈ Θ the set of fixed effects (global)^1^, *c* ∈ 𝒞 the set of covariates, and η ∈ 𝒩 the set of random effects (individual). With slight abuse of notation, constraint (1e) and constraint (1f) restrict fixed and random effects to be drawn from an associated distribution *D*. The subscript *i* denotes the subject. In constraint (1b) the function *g* : Θ × 𝒞 × 𝒩 ↦ 𝒫 maps fixed effects, random effects, and covariates to the set of subject-specific parameters *p*_*i*_ ∈ 𝒫. Constraint (1c) defines the dynamics of each subject. *x* ∈ 𝓍 is a set of dynamical state variables that follow the function *f* : 𝓍 × 𝒰 × 𝒫 × ℝ ↦ 𝓍^′^, often directly dependent on external events *u* ∈ 𝒰 such as drug admissions over time *t* ∈ ℝ. The constraint (1d) describes the observations *y* ∈ 𝒴 using the mapping *h* : 𝓍 × 𝒫 × 𝒞, ×𝒩 × Θ ↦ 𝒴. The objective (1a) of the nonlinear optimization problem (1) is to minimize the negative log-likelihood ℒ : 𝒴 × 𝒴 ↦ ℝ of the observed variables and their estimates by finding the optimal fixed and random effects given a set of data ᵉ𝓍 = 𝒞 ∪ 𝒴 ∪ 𝒰. Typically, optimization problem (1), is solved using a sequential approach (FO) ^2^, as a bilevel optimization problem (FOCE, LaplaceI) ^3^, Expectation Maximization, or fully Bayesian methods, see e.g. [30], [31] or [32] for a general overview.

A common challenge in PKPD and NLME models, but also in general, is to choose the triplet (*g, f*, *h*) so that the structure of the model can represent the variability of the observations using the least number of covariates, fixed and random effects^4^. Especially in the case of an unknown or uncertain mechanism, testing all modeling-driven hypotheses using model selection is a time-, computation- and effort-consuming task. Here, machine learning (ML) offers a data-driven alternative to exhaustive search techniques. Using neural networks (NN) as replacements or extensions of the model can lead to automatically derived functional forms because of the universal approximation capabilities. Essentially, the model itself can formulate a data-driven model hypothesis over the training. A major advantage over classical model selection is the fact that the encoded functional space represented by the NN is smooth in its parameters and, in theory, does not require combinatorial enumeration. In practice, hyperparameter optimization is often needed to find a suitable configuration of the NN in terms of activation functions, number, and width of the layers.

### 2.2 Symbolic Regression

Although neural networks (NN) and other universal approximators are effective, they typically operate as opaque models, offering no insight into the underlying processes beyond the relationships between inputs and outputs. Symbolic regression (SR) provides an alternative approach to techniques that seek to interpret NNs and provide significance to their elements. SR attempts to discover a mathematically parameterized function *f* : 𝓍 × 𝒫 ↦ 𝒴 that accurately represents a set of input and output data *D* = (*x, y*), *x* ∈ 𝓍, *y* ∈ 𝒴 ^5^:

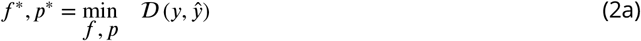

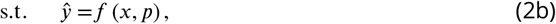

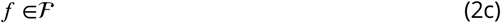

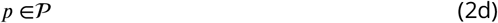

where the unspecified function *f* is utilized to minimize objective (2a) with the mapping ᵉ𝓍 : 𝒴 × 𝒴 ↦ ℝ. The search domain is described abstractly in constraint (2c), typically characterized by its combinatorial nature. Therefore, optimization problem (2) is frequently addressed using metaheuristic approaches such as genetic algorithms, but recently also using reinforcement learning or mixed integer programming [33].

## 3 Methods & Materials

### 3.1 Deep Nonlinear Mixed Effect Models

Deep nonlinear mixed effect (DeepNLME) models enhance traditional NLME models by incorporating neural networks [34]. Within DeepNLME, a neural network is represented by the function 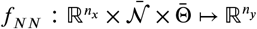, which is characterized by the layerwise computation:

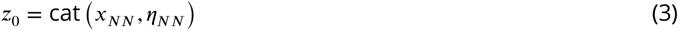

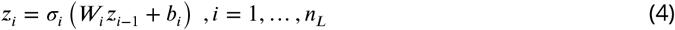

Across all layers, denoted by *n*_*L*_ ∈ ℕ where *n*_*L*_ ≥ 1. Each layer is equipped with an elementwise activation function σ : ℝ ↦ ℝ, weights 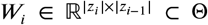, and biases 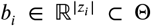, all considered as fixed effects within the model. To handle output variability, a subset of random effects 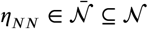 is introduced as an extra input to the neural network (NN) and merged with the input signal 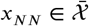.

### 3.2 Typical Value Symbolic Regression

A straightforward method to implement SR within NLME frameworks involves addressing both challenges by extending optimization problem (1) with constraint (2c) and constraint (2d), adding the unknown function directly as an optimization variable to the model. However, particularly when deducing (partially) unknown dynamics, this strategy may result in poor numerical conditioning of the problem. This primarily stems from the exploratory nature of SR, which can inherently cause stiffness, instabilities, or even impractical configurations, making the direct method problematic and the use of smooth universal approximators favorable. Rather, we suggest a sequential fitting method similar to [11]. Initially, we use DeepNLME to train a neural network as a universal approximator to train the missing (sub)model. Subsequently, we employ the derived signals to determine the absent functional form through symbolic regression.

The first stage is effectively carried out with the use of DeepPumas, which diminishes the potential for numerical problems and produces a predictive model that can effectively interpolate across its training data. Moreover, the missing signal 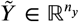 can be directly obtained from the forward pass of the model by simply integrating a new observation, specifically the output of the neural network. Next, our goal is to derive a functional form through SR. Since the trained predictor serves as a substitute for our absent signal, we employ SR as previously mentioned to establish a comprehensive NLME model. This method is generally more adaptable, and we plan to use our trained neural network (NN) as an oracle. Similarly to the FO method, we initially model the network response based on typical values 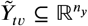 by setting *η*_NN_ as a fixed constant for every input *x* in the population. Subsequently, we apply SR to identify a collection of potential models expressed as equations. We then performed a comprehensive statistical fit using a few of these equations chosen in our original model. This approach is driven by the understanding that output variations should be attributed solely to random effects rather than the model’s expression structure, which is presumed to be invariant across subjects. Therefore, we hypothesize that the basic architecture of the expression can be inferred exclusively from responses based on typical values or an equally suited measure.

### 3.3 Baseline Model

Following [1], we implement a baseline model given by:

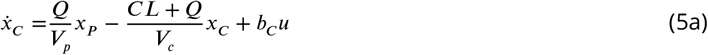

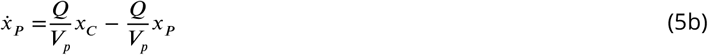

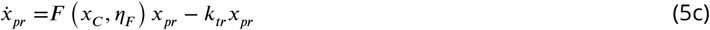

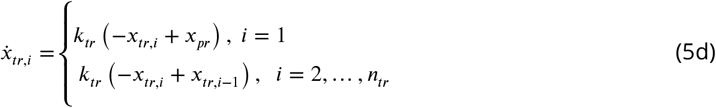

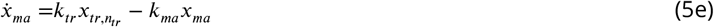

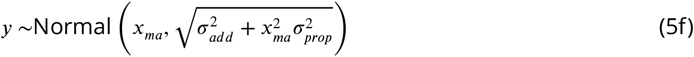

where Equation (5a) and Equation (5b) describe the pharmacokinetic as a central peripheral system with respective states *x*_*C*_, *x*_*p*_, Equation (5c) describes the dynamics of the proliferating cells *x*_*pr*_, Equation (5d) the dynamics of each of the *n*_*tr*_ transition compartments *x*_*tr*_ and Equation (5e) the dynamics of the maturated white blood cells *x*_*ma*_. Equation (5f) describes the observed variable *y* assuming that the WBC is normal distributed with an expected value *x*_*ma*_ using a additive-proportional error model depending on additive and proportional variances σ_*add*_ ∈ ℝ^+^, σ_*prop*_ ∈ ℝ^+^ respectively. The function *F* : 𝓍 × 𝒩 ↦ ℝ couples the effect of the PK and PD models and models the feedback relation :

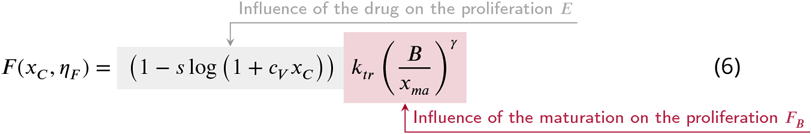

where the influence of *x*_*ma*_ on the proliferation is given as a nonlinear function of the baseline value *B* ∈ ℝ^+^ and the feedback exponent γ ∈ ℝ^+^. The influence of *x*_*C*_ is parameterized by the slope *s* ∈ ℝ^+^ using a constant *c*_*V*_ ∈ ℝ^+^. *E* represents the impact of the drug on the proliferation rate. The feedback term *F*_*B*_ models how mature blood cells affect the growth of new cells. Its overall structure was introduced in [29] and is commonly referred to as the Friberg model. In general, the study and correct modeling of proliferation dynamics is discussed in various sources in the literature, e.g. [35], [36], and various extensions and alternative functional forms are proposed [1], [8].

We choose the initial conditions similiar to [1] ^6^

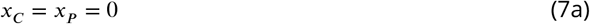

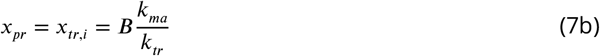

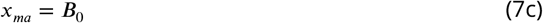

where *B*_0_ ∈ ℝ^+^ is represents the baseline for the WBC in the treatment steady-state of Equation (5). A detailed overview of all parameters, their respective initial conditions, and their description can be found in the supplement Appendix 1.

## 4 Case Study

To investigate our proposed method, we conducted a preliminary study based on publicly available data provided in [1], originally collected from 2008 to 2015 at the Department of Hematology and Oncology, Magdeburg University Hospital, Magdeburg, Germany. It consists of 23 patients who receive high-dose Ara-C infusion induction therapy resulting in complete remission. The white blood cell count (WBC) is measured almost daily. For more details on the subjects, we refer the reader to [1].

### 4.1 Data Preparation

We divided the data into training and validation sets, stratifying based on the average dosing amount and the total observation time relative to the population average. We opted for an 80/20 split, resulting in 18 subjects in the training set and 5 in the validation set. Figure 3 illustrates the WBC count over time and the WBC distribution for both the training and validation sets.

**Figure 3.**
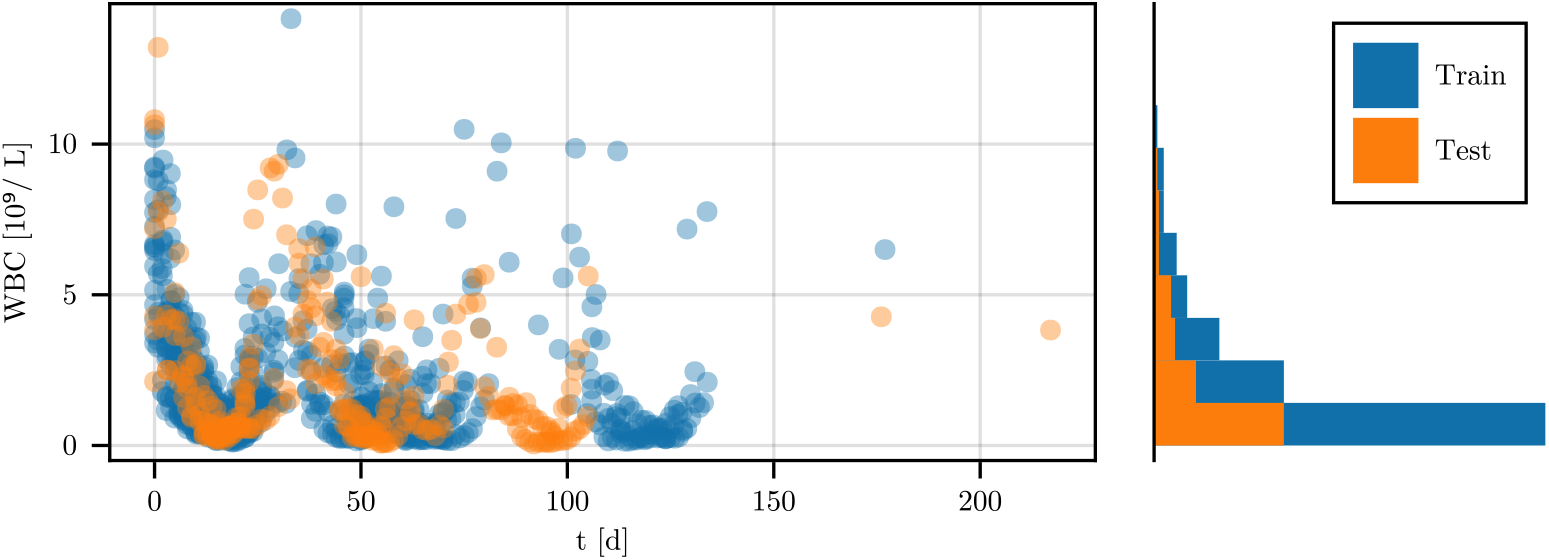
A graphical summary of the provided dataset. **Left:** The WBC count of all subjects over time. **Right:** Histogram of the WBC Count over the study over its number of occurance. Blue denotes the training data, Orange denotes the validation data.

### 4.2 Model Fitting and Evaluation

As an initial step, we repeated the fitting of the PK system as detailed in [1] using the data from [37] with a naive pooled approach, identifying a Central-Peripheral system that achieved the highest loglikelihood value. We created three versions of the baseline model described by Equation (5) by varying the number of transition compartments *n*_*tr*_ = 1, 2, 3. Similarly, we generated ten versions of DeepNLME models where we replaced the feedback function *F*_*B*_ with an ANN for each number of transition compartments, varying the initial distribution of the network’s weights and biases θ_NN_. We used a fixed network structure with three inputs, two hidden layers with seven neurons each, and a single neuron in the output layer. Except for the output layer, tanh : ℝ ↦ {−1, 1} was used as the activation function. The output layer used the softplus function *σ*^+^ : ℝ ↦ ℝ^+^, *σ*^+^ (*x*) = log (1 + exp (*x*)), assuming strictly positive values for the surrogate.

The models were optimized using a maximum a posteriori log-likelihood objective and first-order conditional estimation approximation in Pumas and DeepPumas [38]. We limited the maximum number of iterations to 100 and set a time limit of 6000 seconds. For DeepNLME models, we allowed for increases in the objective function between iterations. As the solver for the dynamical system, we employed an automatic stiffness detection solver using the Vern7 and Rosenbrock23 methods, with absolute and relative tolerances of 1*e* − 6 and 1*e* − 2, respectively.

### 4.3 Symbolic Regression

After training the DeepNLME models, we chose the optimal model for each *n*_*tr*_ based on the loglikelihood of the training set. We then extracted the typical value response 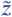 from the neural network and the trajectory data of *x*_*ma*_ for individual predictions of each subject. Using SymbolicRegression.jl [39], we fitted the data with the operator set (+, −, ⋅, ÷, pow), which includes addition, subtraction, multiplication, division, and power. To ensure simplicity, we restricted our search space to a maximum of 12 nodes and prohibited the nesting of the power operator. Each model was fitted for 1000 iterations ^7^ using an *L*_1_ norm objective to create a pareto-frontier of mathematical expressions 𝒰_SR_. Subsequently, we developed a PKPD model for each expression *f*_SR_ ∈ 𝒰_SR_ by substituting *F*_*B*_ and fitting the resulting system as previously described, starting from the optimal values for all fixed effects. Finally, we identified the best model by selecting the one with the lowest BIC on the validation dataset.

## 5 Results

A compact overview of the fitting is given in Table 1. In general, a single transition compartment performed best over all model assumptions, suggesting it as the best option. This is supported by the number of successful results from the DeepNLME model, which shows a decreasing trend with increasing *n*_*tr*_. Given that we directly infer a feedback within the dynamics block of the statistical model having a higher number of transition compartments effectively delays the feedback and hence the model becomes less sensitive to changes in the maturated blood cells, effectively resulting in an instability due to overshoot or undershoot of the equilibria.

**Table 1.** Overview of the results of the fitting procedure. Each baseline and symbolic model has been fitted exactly once, each DeepNLME model is fitted ten times. ℒ denotes the loglikelihood. The number of successful fits and the overall loglikelihood indicate that *n*_*tr*_ = 1 is the optimal structural assumption for the given models.

|  |  | Baseline |  |  | DeepNLME |  |  | Discovered |  |  |
| --- | --- | --- | --- | --- | --- | --- | --- | --- | --- | --- |
| | | | | | $n_{tr}$ | | | | | |
|  | Overall | 1 | 2 | 3 | 1 | 2 | 3 | 1 | 2 | 3 |
| Training $\mathcal{L}$ | | | | | | | | | | |
| Min. | -2635.0 | -705.0 | -735.0 | -992.0 | -2635.0 | -2372.0 | -1985.0 | -655.0 | -702.0 | -735.0 |
| Max. | -639.0 | -705.0 | -735.0 | -992.0 | -646.0 | -659.0 | -663.0 | **-639.0** | -677.0 | -713.0 |
| Mean | -1151.0 | -705.0 | -735.0 | -992.0 | -1394.0 | -1621.0 | -1043.0 | -646.0 | -690.0 | -724.0 |
| Std. | 652 | 0 | 0 | 0 | 799 | 665 | 547 | 6.62 | 18 | 16 |
| Success | 33 | 1 | 1 | 1 | 9 | 7 | 6 | 4 | 2 | 2 |
| Validation $\mathcal{L}$ | | | | | | | | | | |
| Min. | -12991.0 | -252.0 | -258.0 | -412.0 | -1591.0 | -12991.0 | -681.0 | -217.0 | -217.0 | -226.0 |
| Max. | -146.0 | -252.0 | -258.0 | -412.0 | -193.0 | -161.0 | **-146.0** | -214.0 | -214.0 | -222.0 |
| Mean | -876.0 | -252.0 | -258.0 | -412.0 | -677.0 | -2827.0 | -285.0 | -215.0 | -216.0 | -224.0 |
| Std. | 2287 | 0 | 0 | 0 | 552 | 4998 | 231 | 1.14 | 2.09 | 2.96 |
| Success | 31 | 1 | 1 | 1 | 9 | 6 | 5 | 4 | 2 | 2 |
| Training BIC |  |  |  |  |  |  |  |  |  |  |
| Min. | 1397 | 1490 | 1551 | 2065 | 1991 | 2016 | 2024 | **1397** | 1474 | 1533 |
| Max. | 5969 | 1490 | 1551 | 2065 | 5969 | 5443 | 4669 | 1413 | 1499 | 1564 |
| Mean | 2802 | 1490 | 1551 | 2065 | 3487 | 3940 | 2784 | 1407 | 1486 | 1549 |
| Std. | 1464 | 0 | 0 | 0 | 1599 | 1331 | 1094 | 7.73 | 17.1 | 22.5 |
| Validation BIC |  |  |  |  |  |  |  |  |  |  |
| Min. | 510 | 571 | 583 | 892 | 972 | 907 | 877 | **510** | 513 | 530 |
| Max. | 26568 | 571 | 583 | 892 | 3768 | 26568 | 1948 | 541 | 530 | 533 |
| Mean | 2159 | 571 | 583 | 892 | 1940 | 6239 | 1155 | 526 | 521 | 532 |
| Std. | 4632 | 0 | 0 | 0 | 1104 | 9997 | 461 | 12.8 | 11.7 | 2.05 |

**Table 2.** Result of the inference for the baseline model with one transition compartments bootstrapped via five fits of the model. The confidence interval (CI) is choosen to be 0.95. *tv*_*p*_ denotes the typical value of the parameter *p, ω* denotes the prior to the scale of the random effects.

| Parameter | Typical Value | Std. Error. | CI Lower | CI Upper |
| --- | --- | --- | --- | --- |
| $tv_{ktr}$ | 0.20 | 0.02 | 0.18 | 0.23 |
| $tv_{slope}$ | 11.68 | 2.81 | 10.76 | 17.37 |
| $tv_B$ | 5.97 | 0.28 | 5.32 | 5.96 |
| $tv_{B0}$ | 6.45 | 0.12 | 6.32 | 6.59 |
| $tv_{\gamma}$ | 0.48 | 0.09 | 0.40 | 0.62 |
| $\omega_{1,1}$ | 0.05 | 0.01 | 0.04 | 0.06 |
| $\omega_{2,2}$ | 0.42 | 0.18 | 0.23 | 0.67 |
| $\omega_{3,3}$ | 0.15 | 0.04 | 0.10 | 0.21 |
| $\omega_{4,4}$ | 0.01 | 0.00 | 0.01 | 0.01 |
| $\omega_{5,5}$ | 0.43 | 0.12 | 0.16 | 0.42 |
| $\sigma_{prop}$ | 0.38 | 0.03 | 0.35 | 0.42 |
| $\sigma_{add}$ | 0.01 | 0.00 | 0.01 | 0.01 |

**Table 3.** Result of the inference for the discovered model with one transition compartments bootstrapped via five fits of the model. The confidence interval (CI) is choosen to be 0.95. *tv*_*p*_ denotes the typical value of the parameter *p, ω* denotes the prior to the scale of the random effects.

| Parameter | Typical Value | Std. Error. | CI Lower | CI Upper |
| --- | --- | --- | --- | --- |
| $tv_{ktr}$ | 0.19 | 0.02 | 0.15 | 0.20 |
| $tv_{slope}$ | 17.80 | 2.16 | 18.72 | 24.06 |
| $tv_B$ | 5.55 | 0.72 | 4.51 | 6.35 |
| $tv_{B0}$ | 6.31 | 0.43 | 5.77 | 6.79 |
| $tv_{p_1}$ | 1.44 | 0.21 | 0.87 | 1.38 |
| $tv_{p_2}$ | 2.11 | 0.56 | 1.11 | 2.52 |
| $\omega_{1,1}$ | 0.06 | 0.01 | 0.05 | 0.06 |
| $\omega_{2,2}$ | 0.28 | 0.08 | 0.11 | 0.29 |
| $\omega_{3,3}$ | 0.06 | 0.01 | 0.04 | 0.06 |
| $\omega_{4,4}$ | 0.15 | 0.02 | 0.15 | 0.19 |
| $\omega_{5,5}$ | 0.12 | 0.04 | 0.11 | 0.19 |
| $\omega_{6,6}$ | 0.16 | 0.03 | 0.11 | 0.17 |
| $\sigma_{prop}$ | 0.33 | 0.03 | 0.30 | 0.36 |
| $\sigma_{add}$ | 0.09 | 0.03 | 0.09 | 0.15 |

In terms of raw predictive performance measured by the loglikelihood of the individual models over training and validation data, the DeepNLME model was able to outperform both the baseline and the discovered model structure. We conclude that the data set, although small compared to other domains, is able to capture most of the characteristics and train the universal approximator sufficiently to interpolate within the given range of values.

The complexity of the model measured by the BIC in the training and validation data shows that our discovered model is favorable compared to the baseline, although the number of parameters increased ( we neglected γ but added parameters according to the expression found). Despite its high number of parameters, the DeepNLME is still favorable compared to some of the investigated baselines. ^8^

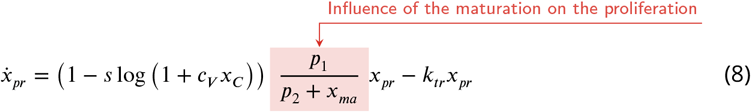

which uses the best expression found via symbolic regression and coincides with the minimum BIC of the validation data according to Table 1. Compared with the Friberg feedback term, our model incoorporates similar characteristics such as inversely proportional behavior but is structurally similar to Hill functions, commonly used to model saturating reactions in pharmacometrics and biology. As such, it can be interpreted as an equilibrium produced by *x*_*pr*_ and *x*_*ma*_ scaled by the impact of *x*_*C*_. Additionally, this expression was found for all DeepNLME instances used as input to SR, strongly suggesting a valid claim for this specific set of data. ^9^ We also choose the best performing models of the baseline (*n*_*tr*_ = 1) and DeepNLME (*n*_*tr*_ = 3). All models produce stable predictions, as can be seen from the individual and typical predictions of Figure 4, Figure 5, and Figure 6. The parity of the fits are comparable, as can be seen from Figure 7, Figure 8, and Figure 9. ^10^

**Figure 4.**
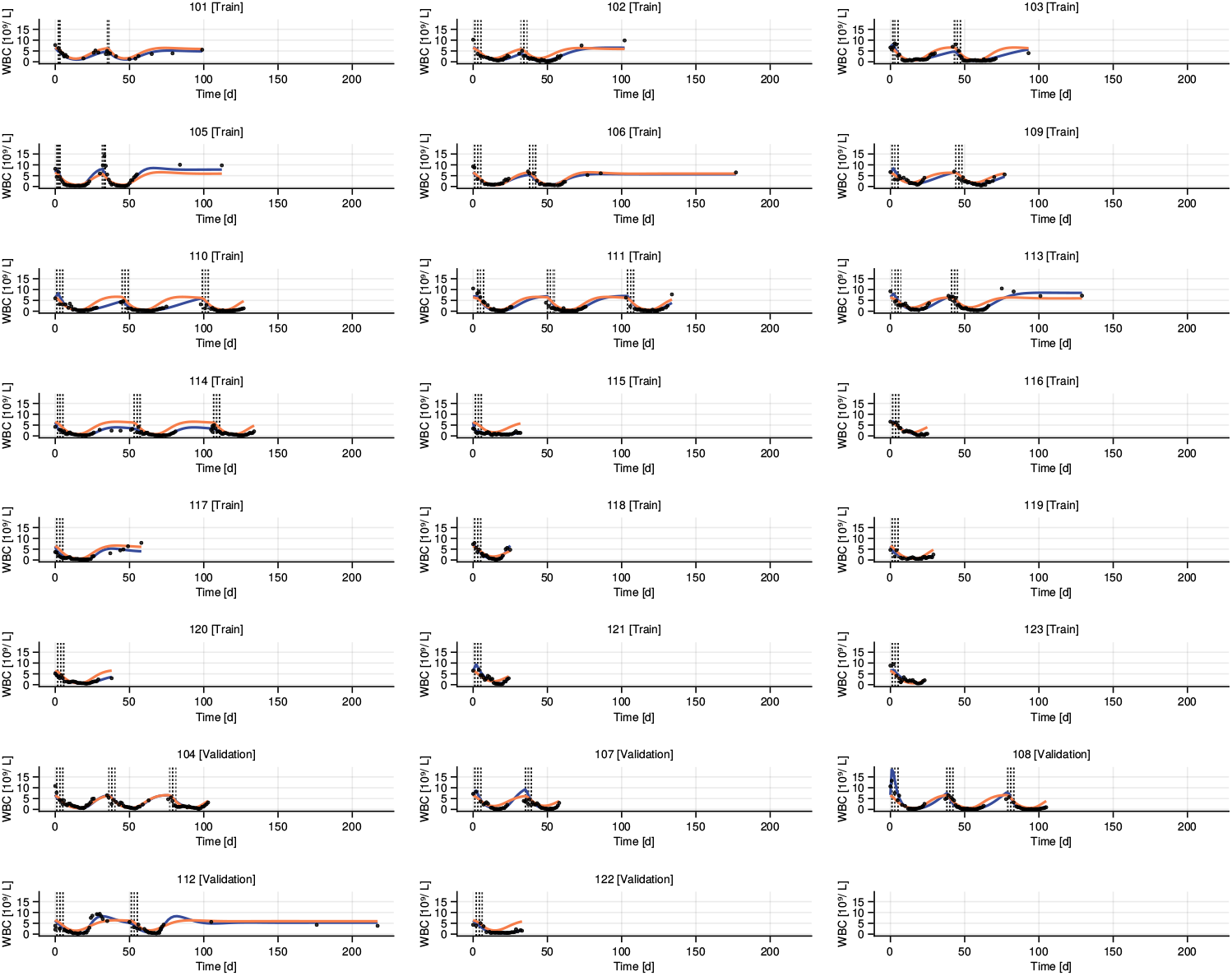
Typical value and individual predictions of the baseline model over both training and validation data.

**Figure 5.**
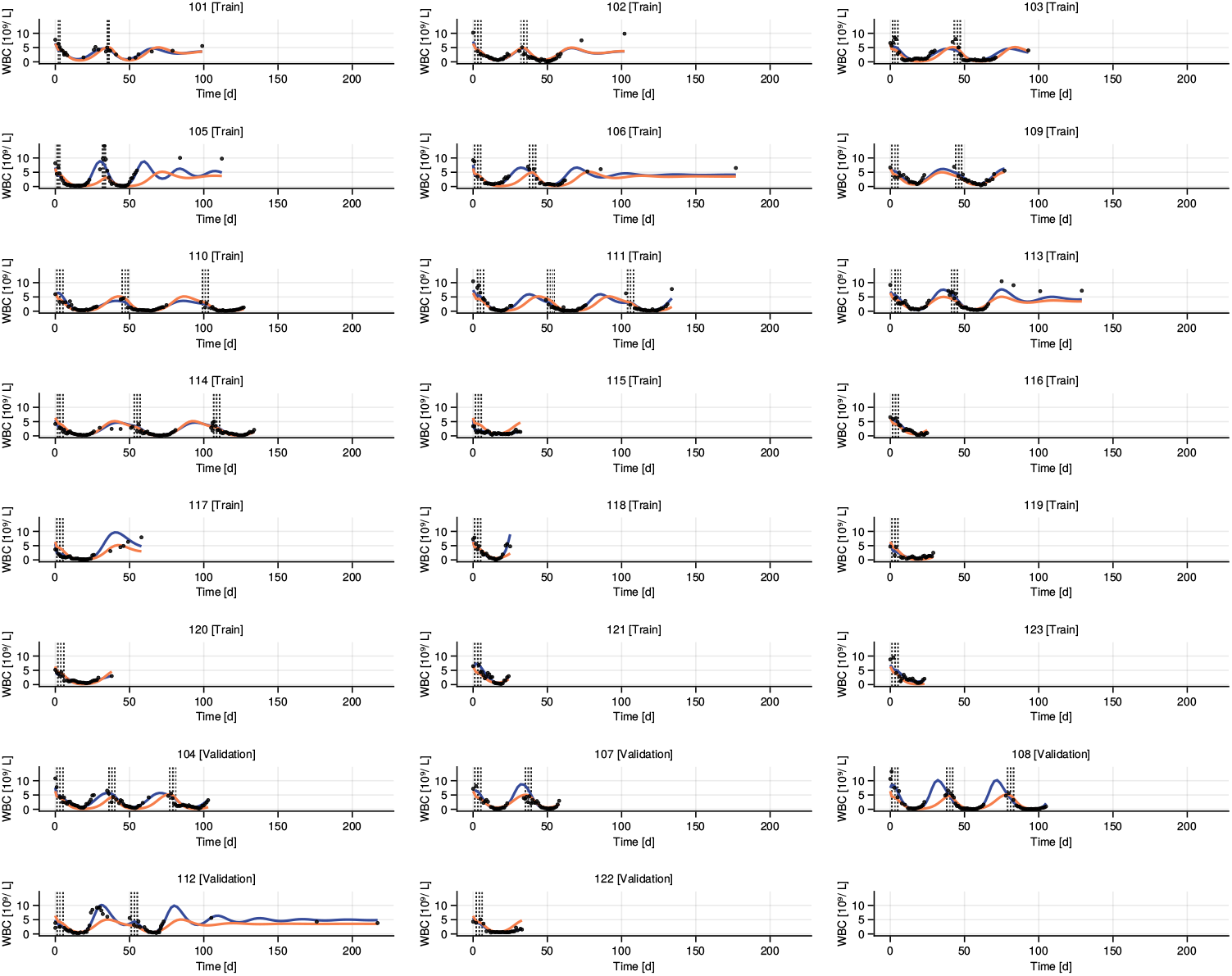
Typical value and individual predictions of the best DeepNLME model over both training and validation data.

**Figure 6.**
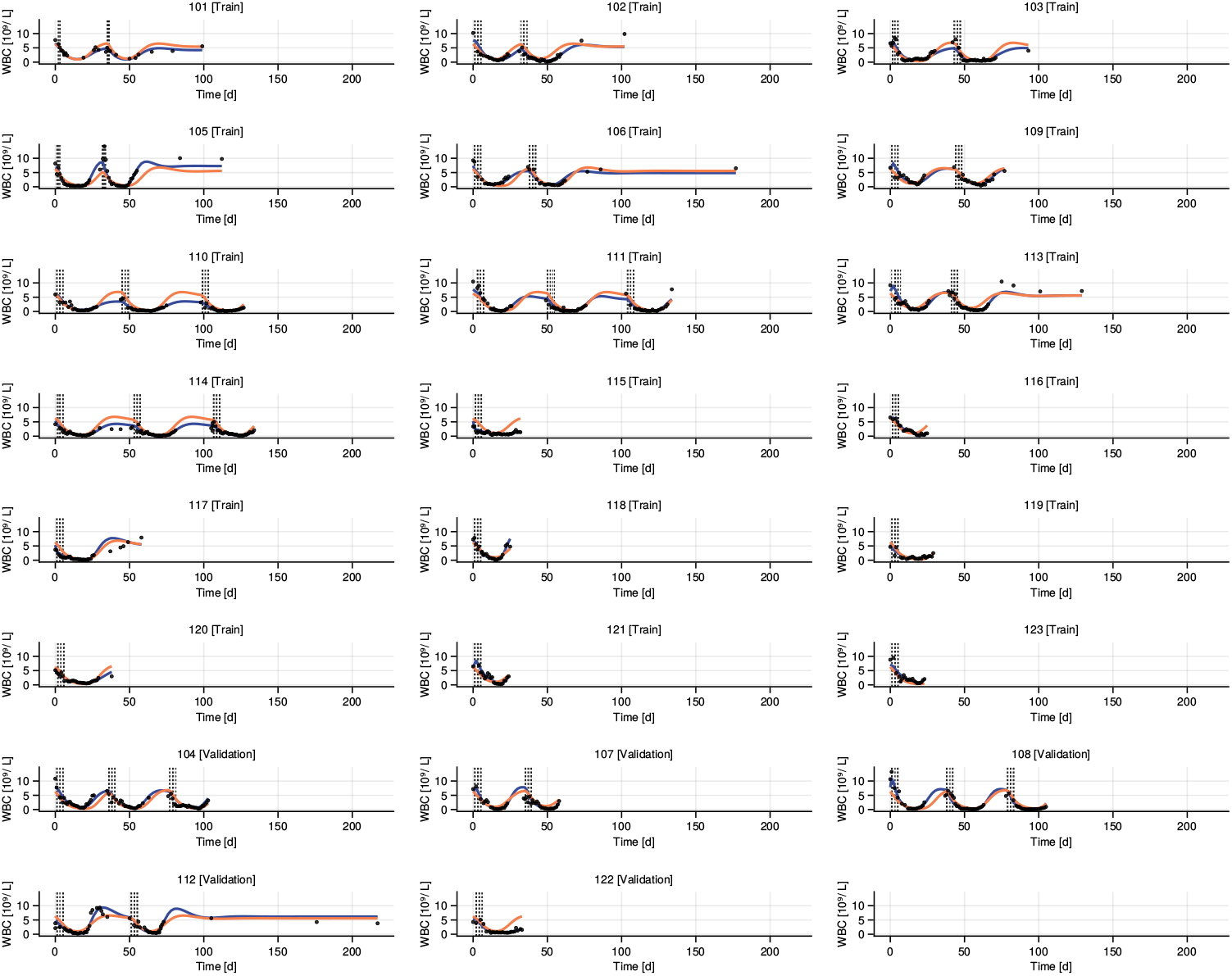
Typical value and individual predictions of the discovered model 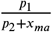 over both training and

**Figure 7.**
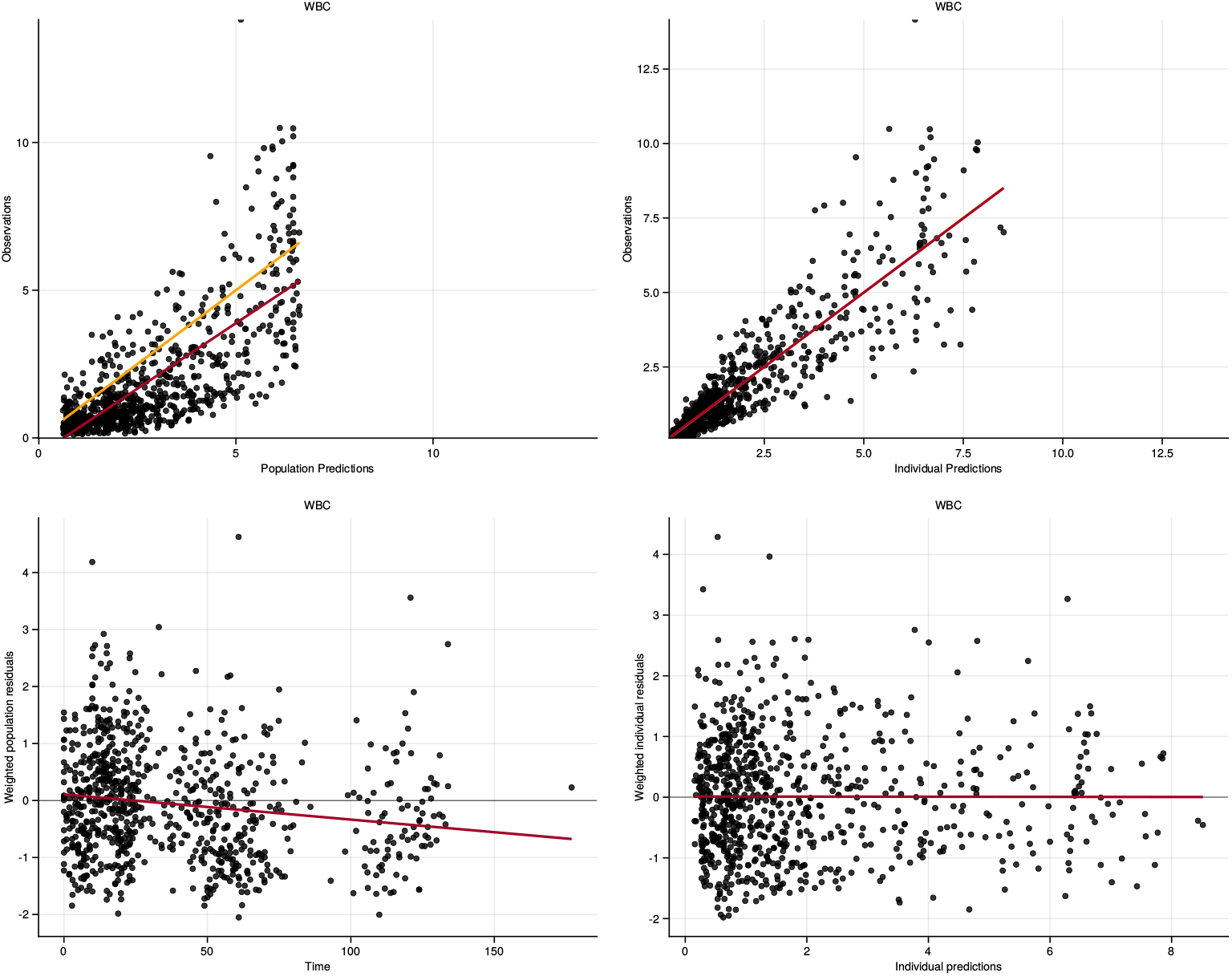
The goodness of fit of the best performing baseline model with *n*_*tr*_ = 1. While the individual predictions capture the data very well a constant offset is present in the typical value predictions. **Upper left:** Parity of the typical value predictions and the observed values. The resulting residuals can be seen in the **lower left. Upper Right:** Parity of the individual predictions and the observed variables. The resulting residuals can be seen in the **lower right**.

**Figure 8.**
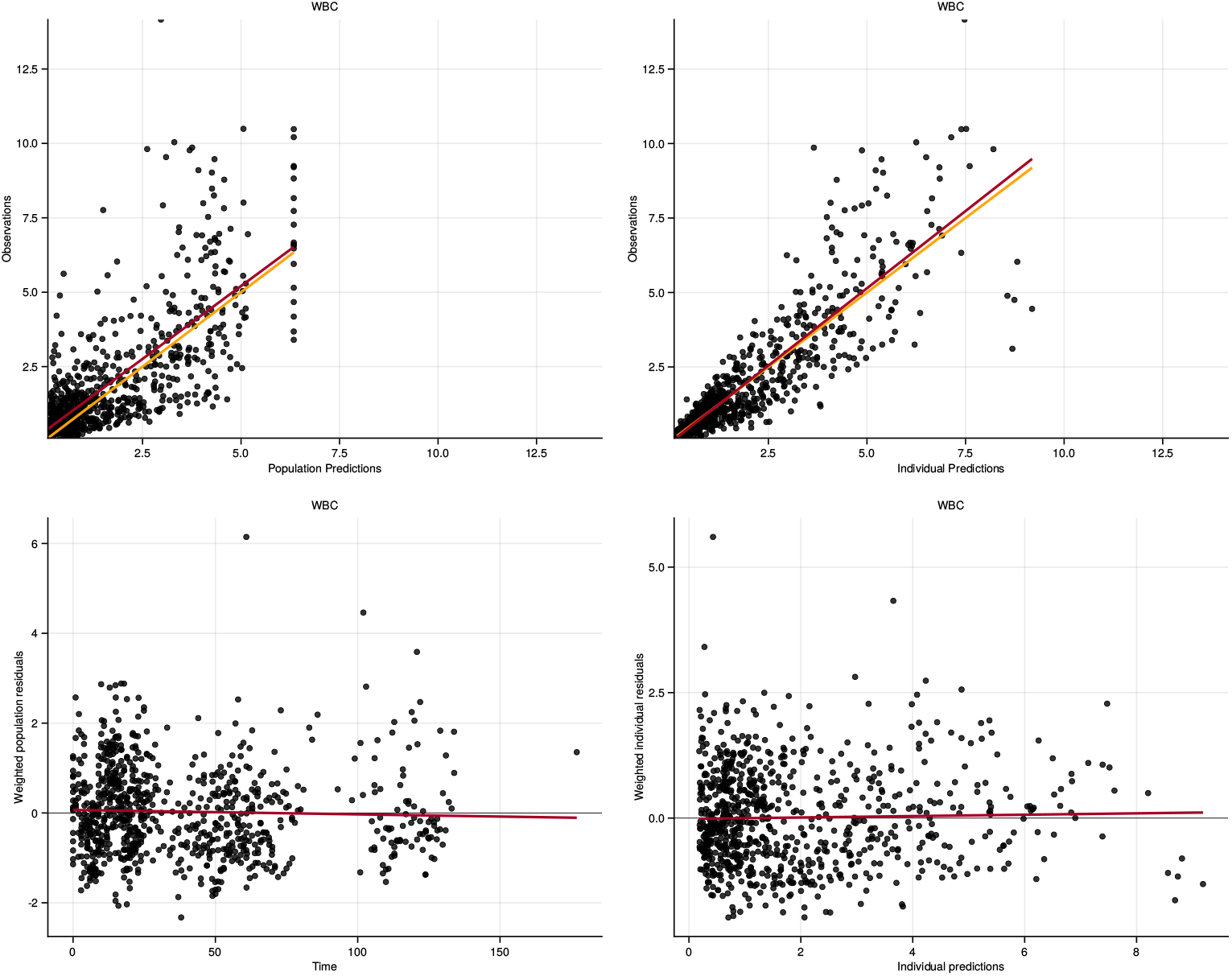
The goodness of fit of the best performing DeepNLME model with *n*_*tr*_ = 3. We can see that both globally and individually the model fits the data almost perfectly due to the universal approximation theorem. **Upper left:** Parity of the typical value predictions and the observed values. The resulting residuals can be seen in the **lower left. Upper Right:** Parity of the individual predictions and the observed variables. The resulting residuals can be seen in the **lower right**.

**Figure 9.**
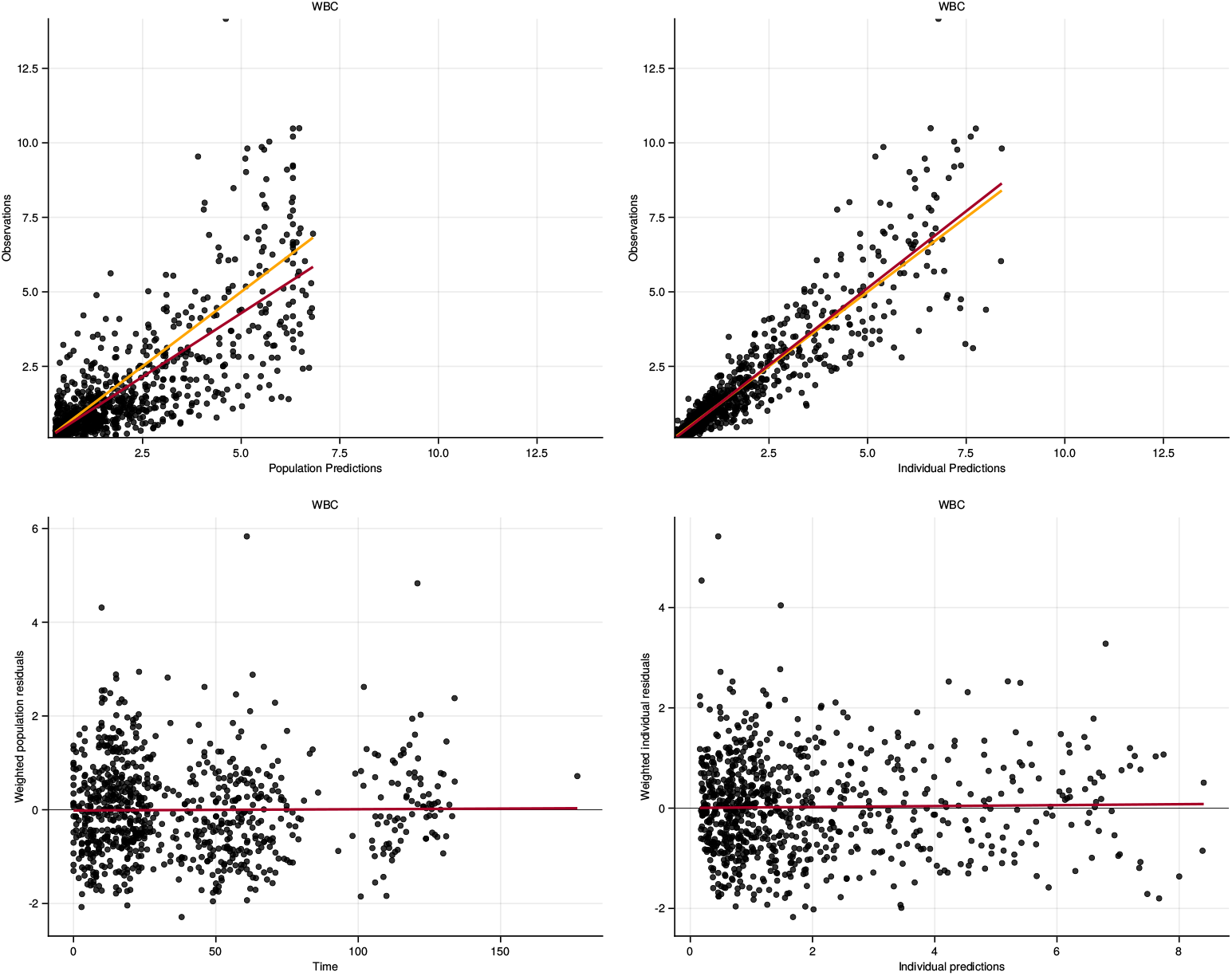
The goodness of fit of the found symbolic model with *n*_*tr*_ = 1. Even though the fit matches the data, it does still incoorporate an increasing offset over the observed variable in both prediction and typical predictions. However, due to its simplicity and numerical robustness it seems favourable over the baseline. **Upper left:** Parity of the typical value predictions and the observed values. The resulting residuals can be seen in the **lower left. Upper Right:** Parity of the individual predictions and the observed variables. The resulting residuals can be seen in the **lower right**.

## 6 Discussion

We presented a method for deriving models directly from data within the framework of nonlinear mixed effect modeling. This method relies on a data-driven surrogate, such as DeepNLME. The case study provided illustrates that the proposed approach to symbolic regression in statistical modeling can discover expressions even in highly nested nonlinear models with external inputs, as is typical in PKPD modeling. Our model retains all the features of the original Friberg function but shows greater resemblance to the widely used class of Hill functions. Additionally, we showed that while DeepNLME models are less interpretable than symbolic expressions, their overall predictive performance on both training and validation datasets supports their use in practical applications. We hope this study marks the beginning of generalized symbolic regression and a stronger emphasis on data-driven modeling not only in PKPD models but in statistical modeling as a whole.

## 6.1 Acknowledgment

This project has received funding from the Deutsche Forschungsgemeinschaft (DFG, German Research Foundation) (No. 314838170), grants SPP 2331 and GRK 2297, from the European Regional Development Fund (grant timingMatters) under the European Union’s Horizon Europe Research and Innovation Program, and was supported by a research grant from the International Max Planck Research School (IMPRS) for Advanced Methods in Process and System Engineering in Magdeburg, which we gratefully acknowledge.

Carl Julius Martensen has been working as an intern at Pumas-AI Inc. during the time of this research. Niklas Korsbo, and Vijay Ivaturi are employed by Pumas-AI Inc.

The authors thank Mohamed Tarek, Andreu Vall, Lorenzo Contento, and Adrian Manuel Reimann for their feedback and insights throughout this study.

This preprint was created using the LaPreprint template (https://github.com/roaldarbol/lapreprint) by Mikkel Roald-Arbøl.

## 6.2 Supplementary

Insert the supplementary text here.

## Appendix 1

### A Initial Conditions

**Appendix 1—table 1.**
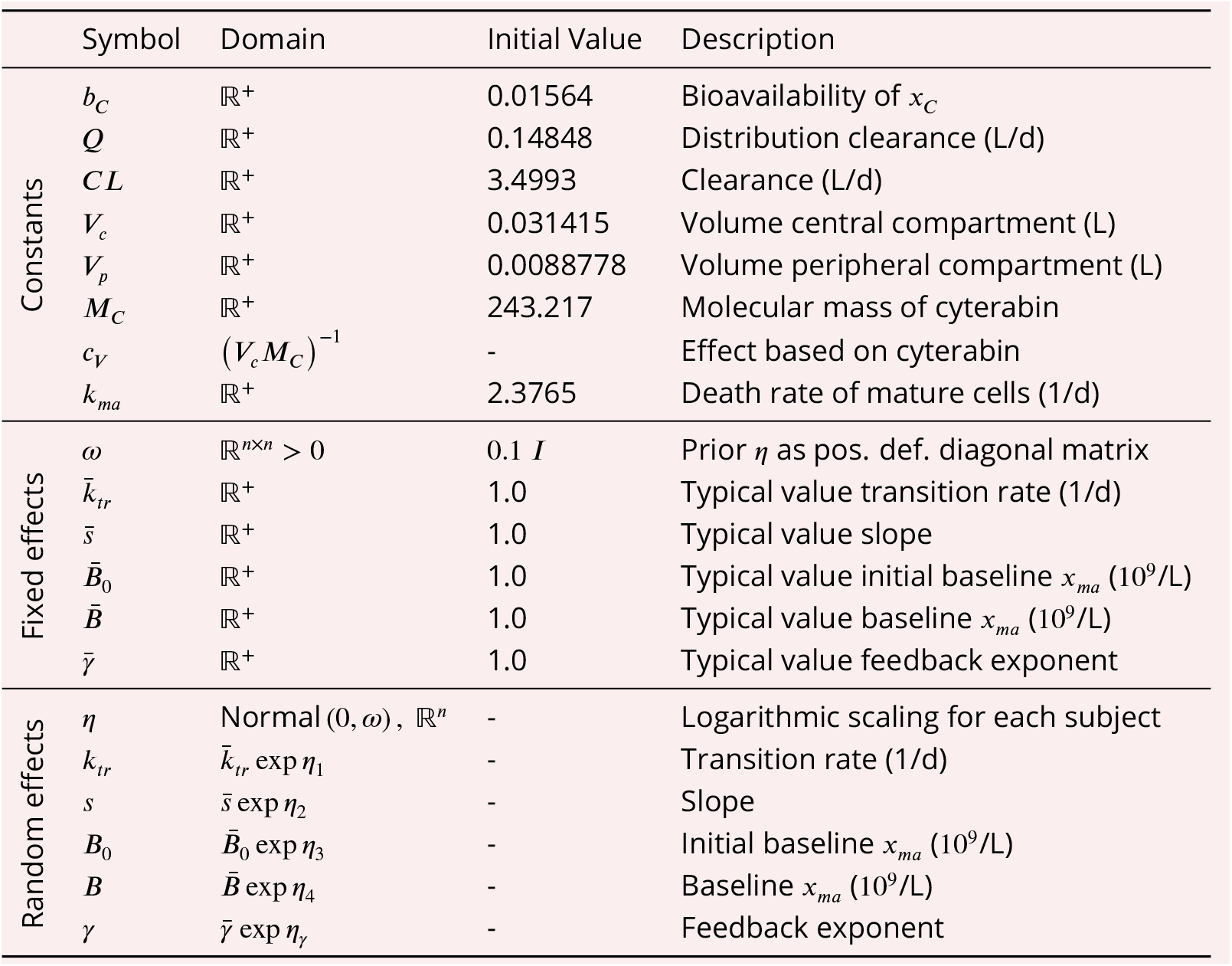
Constants, fixed and random effects and their respective domain, initial value, and description.

## Appendix 2

### B Additional Optimization Information

#### B.1 Visual Predictive Checks

**Appendix 2—figure 1.**
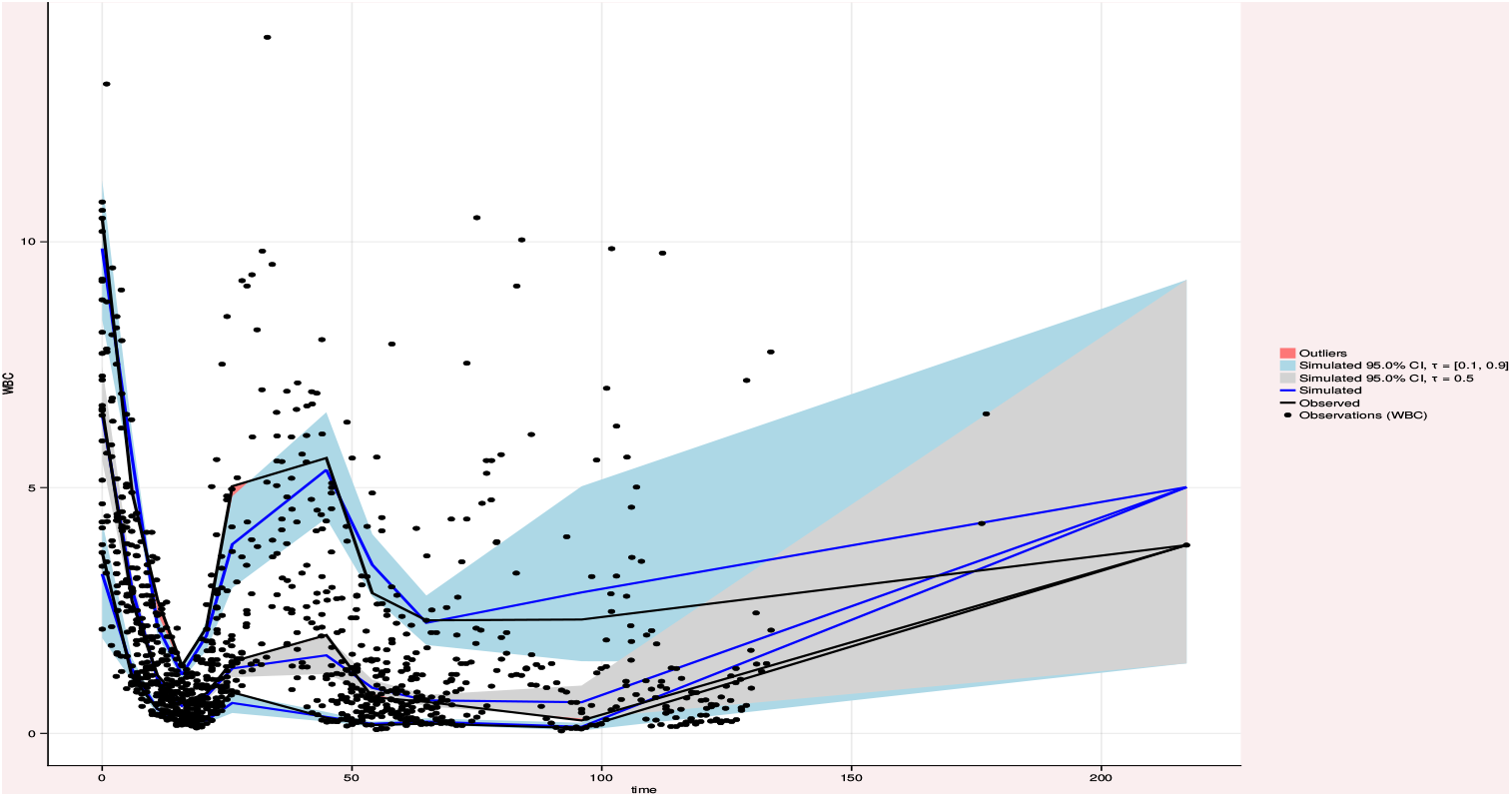
Visual predictive check of the best baseline model.

**Appendix 2—figure 2.**
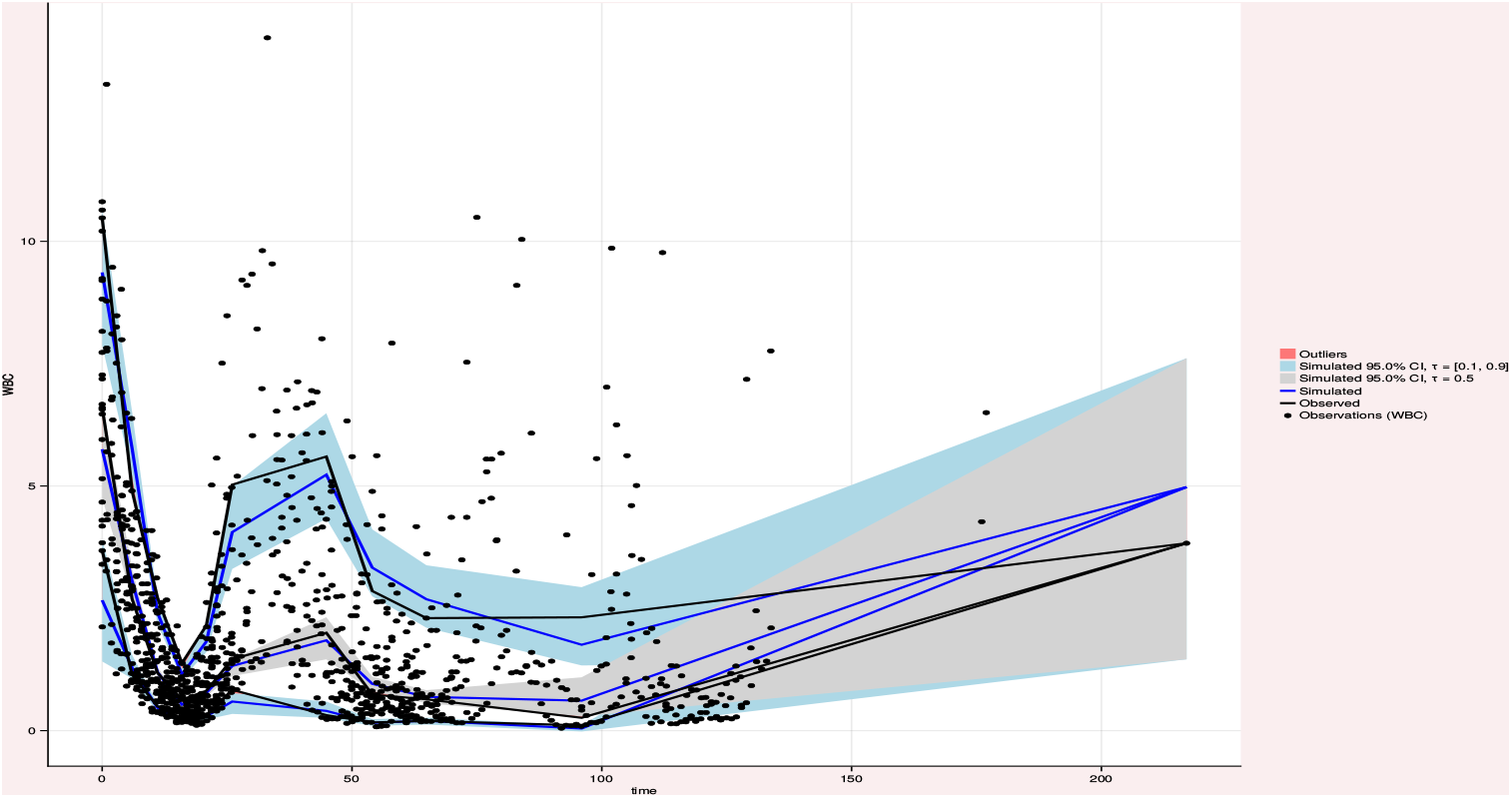
Visual predictive check of the best DeepNLME model.

**Appendix 2—figure 3.**
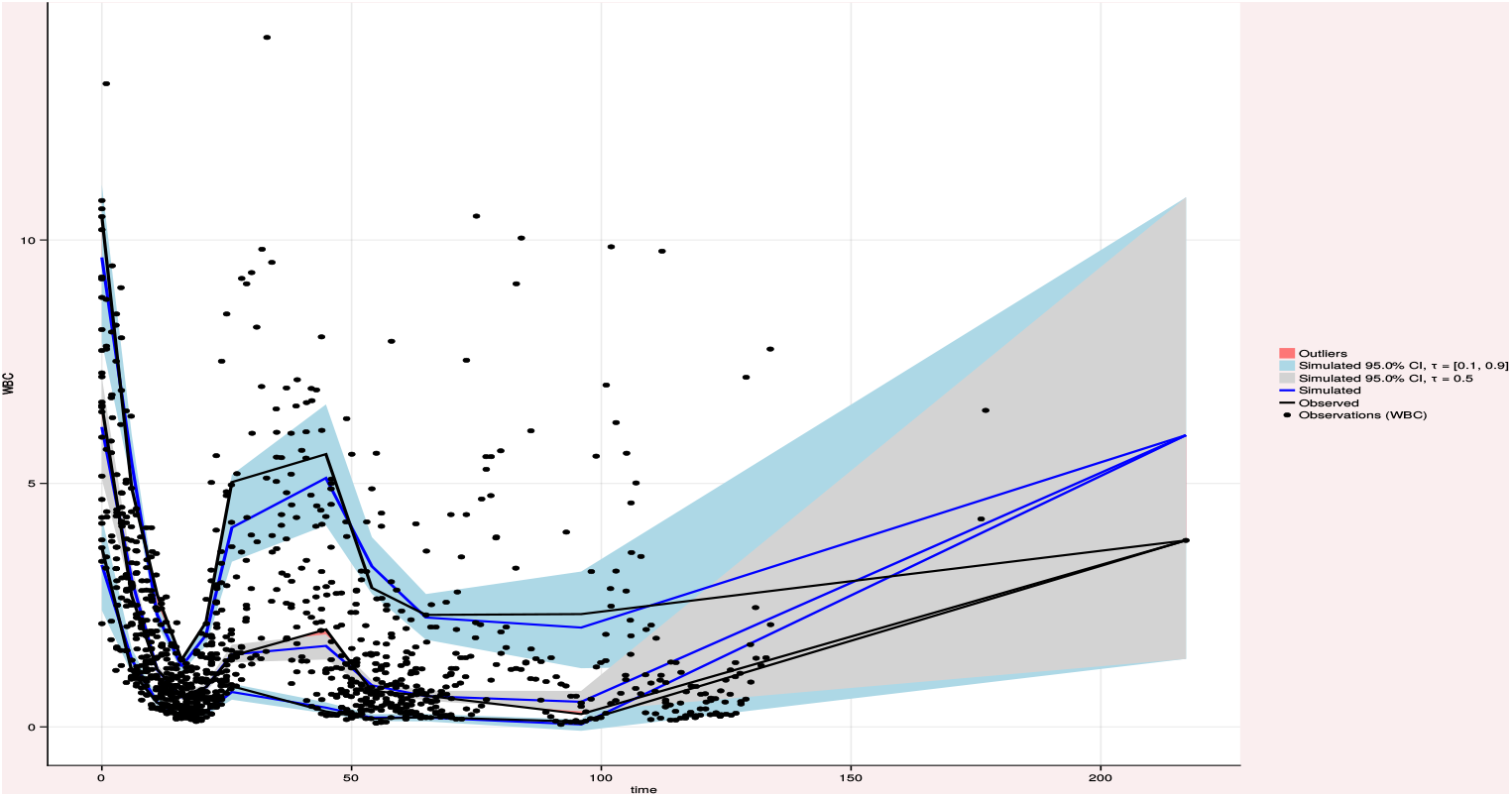
Visual predictive check of the best disovered model.

#### B.2 Empirical Bayes Distribution

**Appendix 2—figure 4.**
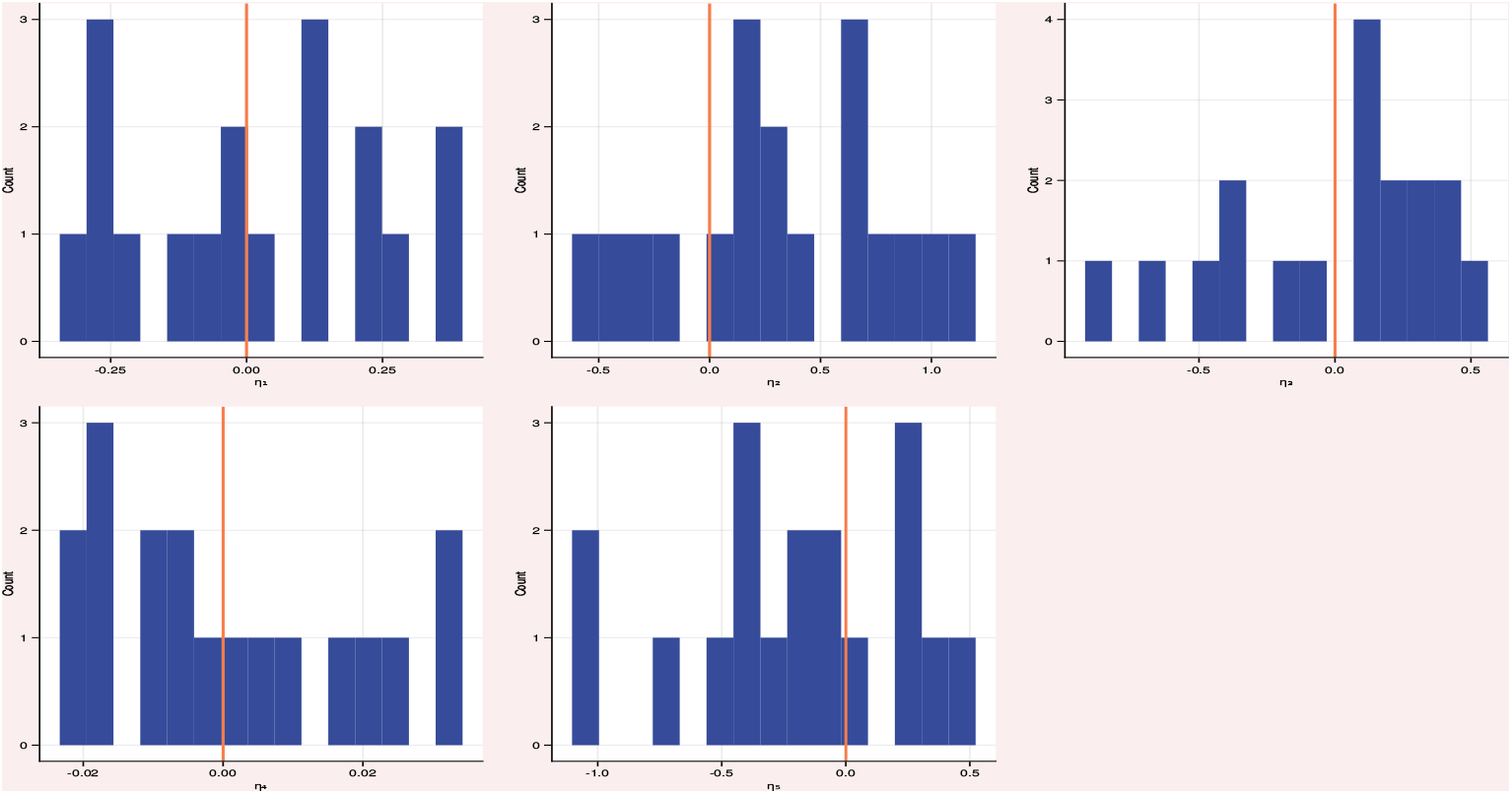
Distribution of the random effects *η* for the best baseline model.

**Appendix 2—figure 5.**
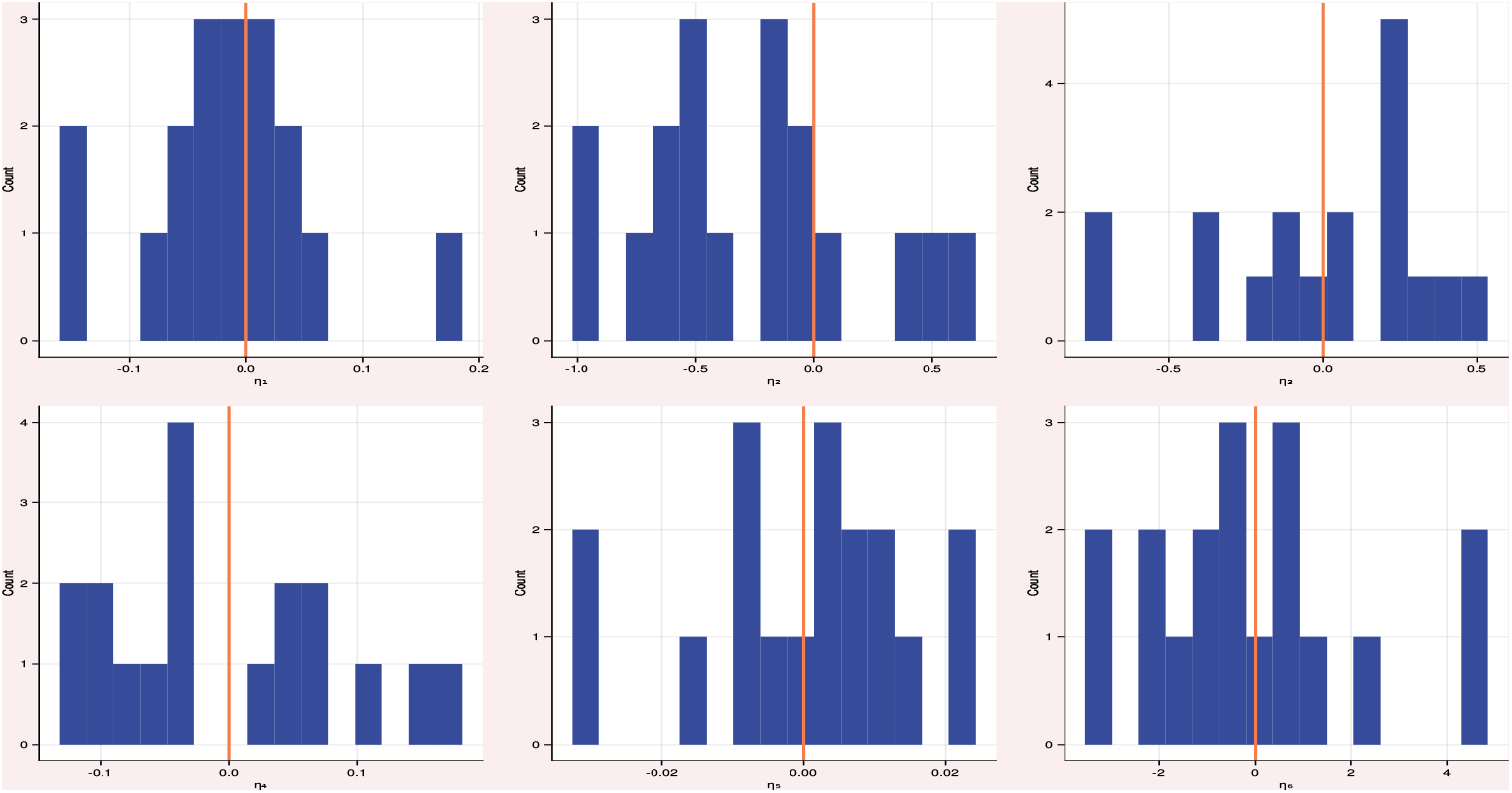
Distribution of the random effects *η* for the best DeepNLME model.

**Appendix 2—figure 6.**
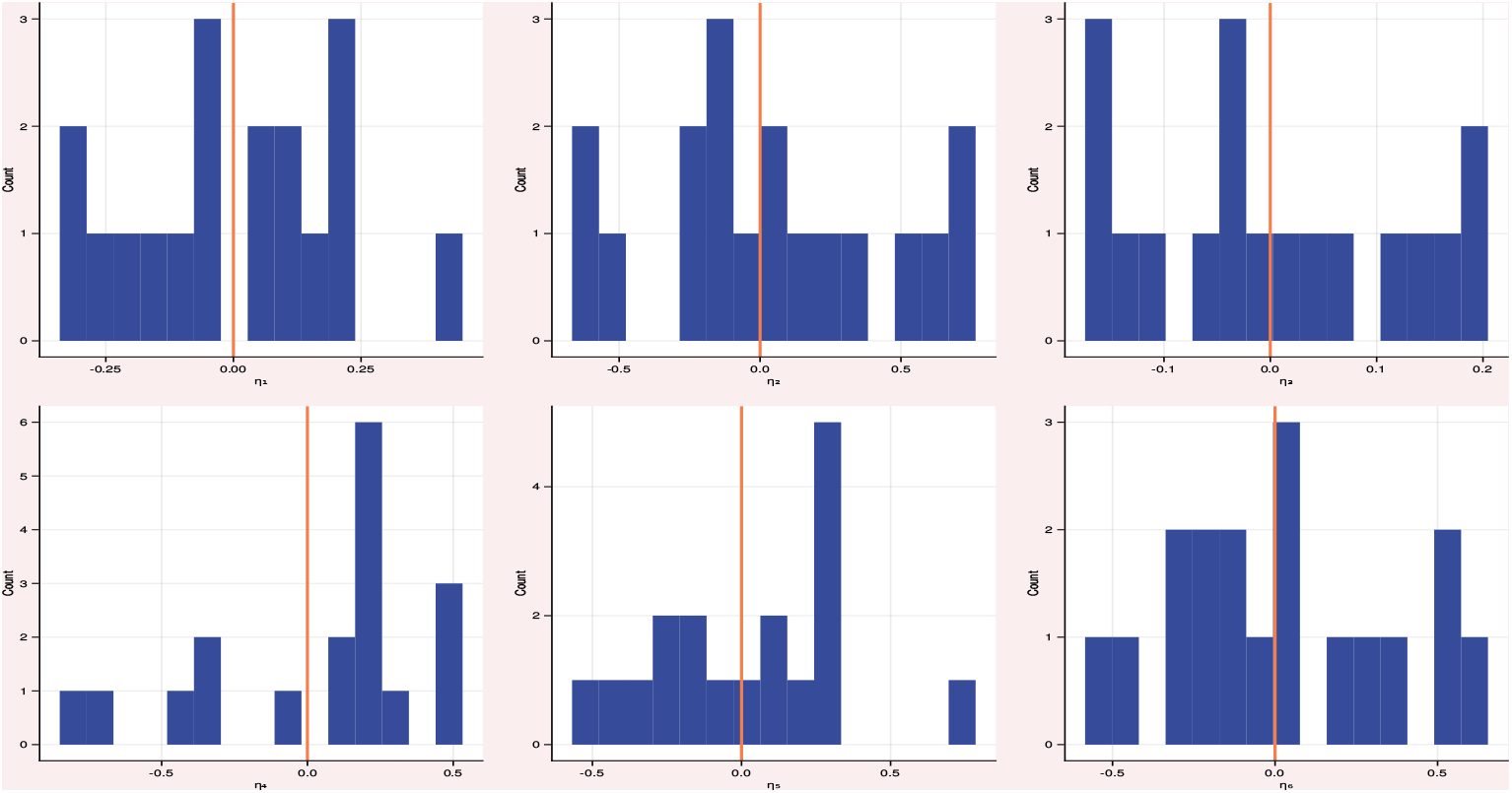
Distribution of the random effects *η* for the best discovered model.

## Appendix 3

### C Symbolic Regression Results

Table 1 shows all expressions found by our method which resulted in a successful fit. In general the results of the symbolic regression where identical up to the values of the parameters.

**Appendix 3—table 1.**
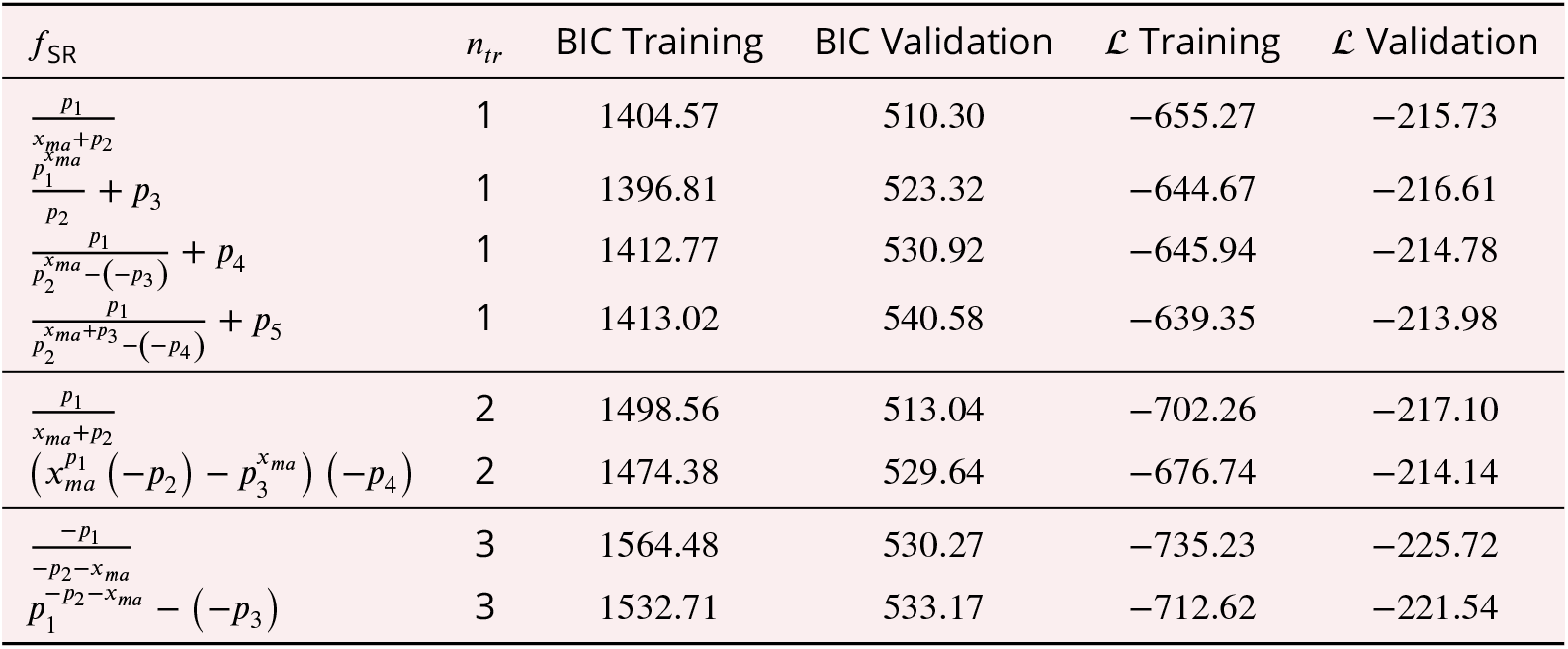
A tabular summary of all expressions identified in the top-performing DeepNLME models for each *n*_*tr*_ that led to successful fits.

## Footnotes

1 We use **global** to refer to parameters which are valid for all subjects and **individual** for parameters which are only valid for a single subject.

2 Optimize *θ* first, then *η*.

3 Optimize *θ* on the outer level, *η* on the inner level.

4 Typically refered to as Occam’s razor or model parsimony. Common measures are the Aikakes information criterion or the Bayesian information criterion.

5 The variables *f, x, y, p* and their respective sets are not related to the variables introduced in optimization problem (1) but are choosen for reasons of brevity and to match the commonly used notation *y* = *f*(*x, p*)

6 This reflects the **treatment steady-state** of the system. Unlike [1], we effectively choose the model M4 with initial conditions I2.

7 Typically, more iterations are required, but we found that the discovered functions usually converged within the first 100 iterations.

8 This could point to a structural error in the modeling approach, the initial conditions, or in the implementation, but does not affect the results of this study given that we assume our model is correct and investigate if DeepNLME and SR can produce better alternatives.

9 A complete list of the expressions derived from the data can be found in Appendix 3.

10 Additional plots can be found in Appendix 2.

## Notes

### Competing Interest Statement

C.J.M., N.K., and V.I. are associated with Pumas-AI.

